# Evaluating strategies for reversing CRISPR-Cas9 gene drives

**DOI:** 10.1101/144097

**Authors:** Michael R. Vella, Christian E. Gunning, Alun L. Lloyd, Fred Gould

**Affiliations:** North Carolina State University, Biomathematics Graduate Program, Department of Mathematics, Raleigh, 27695, USA; North Carolina State University, Genetic Engineering and Society Center, Raleigh, 27695, USA; North Carolina State University, Department of Entomology and Plant Pathology, Raleigh, 27695, USA

## Abstract

A gene drive biases inheritance of a gene so that it increases in frequency within a population even when the gene confers no fitness benefit. There has been renewed interest in environmental releases of engineered gene drives due to recent proof of principle experiments with the CRISPR-Cas9 system as a drive mechanism. Release of modified organisms, however, is controversial, especially when the drive mechanism could theoretically alter all individuals of a species. Thus, it is desirable to have countermeasures to reverse a drive if a problem arises. Several genetic mechanisms for limiting or eliminating gene drives have been proposed and/or developed, including synthetic resistance, reversal drives, and immunizing reversal drives. While predictions about efficacy of these mechanisms have been optimistic, we lack detailed analyses of their expected dynamics. We develop a discrete time model for population genetics of a drive and proposed genetic countermeasures. Efficacy of drive reversal varies between countermeasures. For some parameter values, the model predicts unexpected behavior including polymorphic equilibria and oscillatory dynamics. The timing and number of released individuals containing a genetic countermeasure can substantially impact outcomes. The choice among countermeasures by researchers and regulators will depend on specific goals and population parameters of target populations.

## Introduction

Recent work has employed the CRISPR-Cas9 system[8, 14] to create homing drives (HD) that increase the frequency of genetic constructs in a population even if they lower the fitness of individuals that carry them[11]. The drive mechanism exploits homology directed repair (HDR) to replace a targeted, naturally occurring genomic sequence with an engineered construct[4, 10]. The HD construct codes for Cas9 (or any similar endonuclease, such as Cpf1[24]) and one or more guide RNAs so that in HD heterozygotes, the combined presence of Cas9 and the guide RNA(s) converts germline cells into HD homozygotes. The engineered construct may also include a novel, expressed gene.

An HD can be used in two different ways: for population suppression (suppression HD), where the drive induces a major genetic load[2], or for population replacement (replacement HD), where the expressed gene in the drive construct induces an intended phenotypic alteration, such as blocked transmission of a pathogen[11]. Despite their promise, HDs carry a number of potential risks, including unforeseen ecological consequences and unintended geographical spread[10, 17]. The severity of adverse effects could vary widely, with HD individuals and Cas9 remaining in the population. For example, the magnitude of such adverse impacts would likely be affected by the likelihood of undesirable HD migration and by the likelihood of low-probability events such as horizontal gene flow. In some instances, actions to gradually reduce HD frequency may be viewed as sufficient, while in other cases, swift, complete elimination of HD and restoration of the wild-type would be preferred.

It is possible for an HD bearing a fitness cost to naturally go extinct due to evolution against it, such as the spread of drive-resistant alleles developed via non-homologous end joining (NHEJ)[1, 2, 22]. This would likely prevent the HD from reaching fixation but not reduce HD frequencies quickly. HD constructs could also be engineered (i.e., no pre-existing resistant alleles in the population, and multiple guide RNAs to force simultaneous events of NHEJ for resistant alleles to arise) to minimize the likelihood of natural resistance[6, 10]. Thus, several countermeasures have been proposed to proactively slow the spread of an HD and/or remove it from a population. In the case of a suppression HD, one option would be to release individuals carrying a synthetic allele of the targeted gene that is resistant to the HD[2, 6]. In this case, the synthetic resistant (SR) allele would have no substantial fitness advantage over a replacement HD designed to have minimal fitness cost.

A second option that could be useful for stopping either suppression or replacement HDs involves synthetic CRISPR-Cas9 based overwriting or reversal drives (RD)[7, 10]. CATCHA (Cas9-triggered chain ablation) and ERACR (elements for reversing the autocatalytic chain reaction) have been proposed as RDs[12, 23]. The CATCHA and ERACR constructs contain guide RNAs but do not include the Cas9 gene, depending instead on Cas9 present from the HD. The guide RNAs produced by the RD target the HD construct in the same way that the HD targets the wild-type allele. A third option is using an immunizing reversal drive (IRD) that would target both HD and wild-type populations by including both the Cas9 gene and multiple guide RNAs that target the HD and wild-type sequences[10]. IRDs are designed to replace both HD-bearing and wild-type individuals, with constructs that have active Cas9 and guide RNA production but no intended effect on the organisms phenotype.

The National Academies of Sciences, Engineering, and Medicine report[17] recommended the use of mathematical models in evaluating strategies for reducing potential harms of gene drives. An intuitively reasonable expectation, for example, is that RDs could be employed to eliminate an HD[12]. Yet there has been no quantitative assessment to date of the predicted dynamics of reversal and immunizing drives. Here we present a simple, frequency-only population genetics model to elucidate the evolutionary dynamics of genetic strategies for countering HDs. We show that SR alleles and RDs are not guaranteed to eliminate an HD from a population due to the existence, in general, of a stable polymorphic equilibrium in which the countermeasure co-exists with the wild-type and HD. An IRD, on the other hand, is much more likely to eliminate an HD but is also expected to eliminate wildtype alleles and continue production of Cas9.

## Methods

We build on previous deterministic models of HD allelic dynamics that employ non-overlapping generations (i.e., a discrete-time description) and random mating[6, 22]. We add alleles for SR, RD, and IRD as countermeasures. Alleles for natural resistance are also examined. We assume that Cas9 always produces a double-strand break in wild-type/HD, HD/RD, wild-type/IRD, and HD/IRD heterozygotes. We assume that resistant alleles arise naturally (and only) via NHEJ whenever HDR is unsuccessful, such that the homing rate is equivalent to the probability of HDR. Finally, we assume that fitness costs yield an excess of lethality relative to wild-type at some point prior to reproduction, and that Cas9 is produced only in the germline. Note that, due to drive activity, gamete genotype contribution may differ, but conversion occurs only after somatic mortality via fitness cost is assessed.

We let *q*_*W*_, *q*_*HD*_, *q*_*C*_, and *q*_*R*_ be the current generation frequencies of wild-type (*W*), HD, countermeasure (*C*), and naturally resistant (*R*) alleles in the population, respectively. The equations predicting the next generation frequencies (*q*′) are:

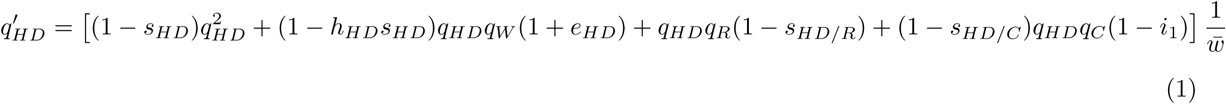

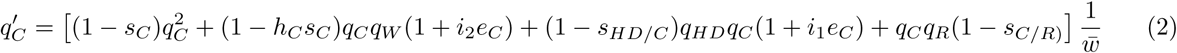

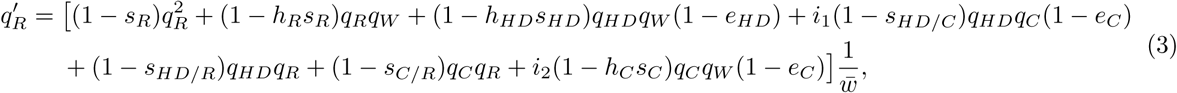

where *q*_*W*_ = 1 − *q*_*HD*_ − *q*_*C*_ − *q*_*R*_ because the frequencies must add to one. The mean population fitness 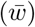 can be calculated by subtracting fitness cost deaths from one:

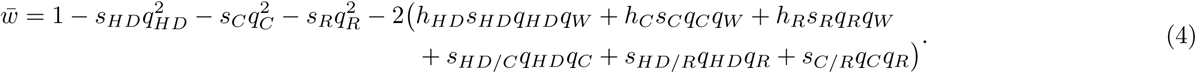

Parameters *e*_*HD*_ and *e*_*C*_ are the probabilities of successful copying (homing) for the homing drive and counter-measure, respectively. The countermeasure allele represents SR when *i*_1_ = *i*_2_ = 0 (no homing), an RD when *i*_1_ = 1 and *i*_2_ = 0 (homing only in HD/countermeasure heterozygotes), and an IRD when *i*_1_ = *i*_2_ = 1 (homing in both HD/countermeasure and wild-type/countermeasure individuals).

We assume wild-type fitness is 1, and define *s* to be the fitness cost of homozygotes. The degree of dominance, *h*, gives the fraction of the homozygote fitness cost imposed on a heterozygote with one wild-type allele. We denote fitness costs of HD/R, HD/C, and C/R heterozygotes as *s*_*HD/R*_, *s*_*HD/C*_, and *s*_*C/R*_, respectively. We assume fitness costs are recessive, with heterozygotes bearing the lesser fitness cost of its alleles, unless noted otherwise. Note that the RD and IRD may recode for the gene interrupted by the HD or eliminate an expressed gene in the HD construct such that the countermeasure constructs do not carry the same fitness costs as the HD construct.

## Results

In Figure 1, we show several examples of countermeasure dynamics that are indicative of behavior over a broad range of parameter values. In these examples, the countermeasures are deployed against a suppression HD, and we assume perfect homing. Figure 1a shows the rapid spread of the HD in the absence of countermeasures, where high HD fitness costs would result in population suppression or extinction. Figure 1, b-g, compares impacts of release of an SR allele (Fig. 1b-c); release of an RD (Fig. 1d-e); and release of an IRD (Fig. 1f-g), with each initiated using a single release of either a 1:1 (Fig. 1b/d/f) or a 1:10 (Fig. 1c/e/g) ratio into populations at the end of the 8th generation after the HD release, when the HD frequency has exceeded 0.2. Regardless of release size, the systems with SR and RD releases reach stable, polymorphic equilibria in the long term, whereas the IRD eliminates the HD and reaches fixation. The SR (Fig. 1b-c) reaches high frequencies and slowly diminishes HD frequencies, though ongoing conversion of wild-type to HD is sufficient to maintain the HD in the population. The larger release of RD (Fig. 1d) immediately brings the system close to the equilibrium, causing HD frequencies to stay relatively constant. The smaller RD release (Fig. 1e) allows HD frequencies to initially increase, which may not be desirable. However, the subsequent buildup of RD then reduces HD to very low frequencies, in contrast to what was seen in Figure 1d. In this trough of low HD frequency, stochastic loss of HD via drift may occur, with the HD loss probability increasing as population size decreases[13]. The IRD does not coexist with other alleles because it maintains an advantage over each of the other alleles regardless of its frequency and quickly reaches fixation regardless of release size (Fig. 1f-g).

Moving to consider replacement HDs, Figure 2 shows a set of time series for an HD with lower fitness cost (*s*_*HD*_), but with otherwise identical parameter values as shown in Figure 1. The qualitative behavior in the replacement HD setting is similar to the behavior in the suppression HD setting, but the lower HD fitness cost slows dynamics. The difference is most notable for the RD with a small release of countermeasure, for which the system exhibits large, slowly damped oscillations that bring the target HD to low frequencies for many generations (Fig. 2e). Due to genetic drift, the likelihood of stochastic loss of an allele increases as the time spent with few copies of that allele in the population increases[13].

**Figure 1:**
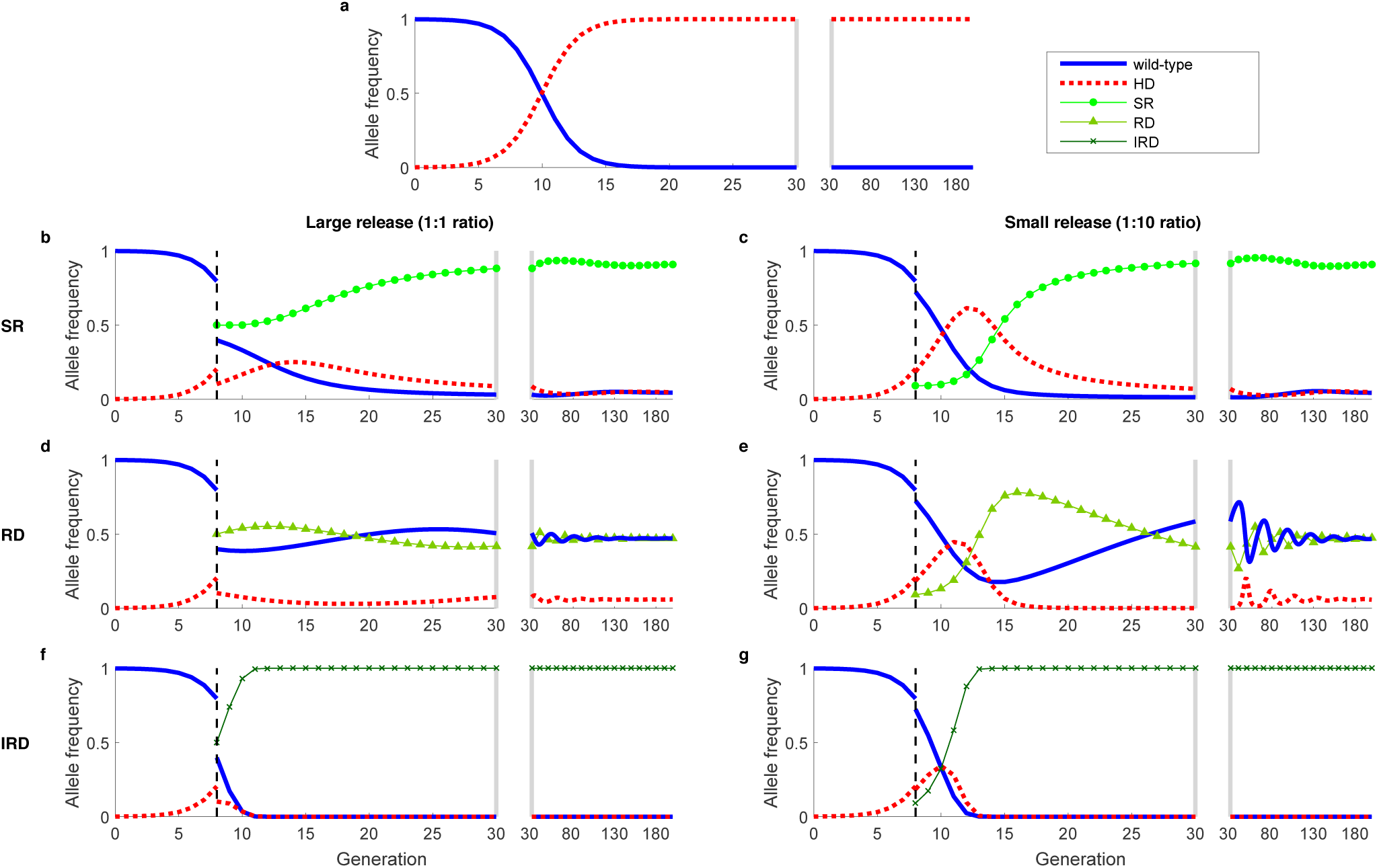
**Dynamics of a suppression HD, alone (a) and with countermeasures (b-g)**, which include a synthetic resistant allele (b,c), reversal drive (d,e), and immunizing reversal drive (f,g). Fitness cost (*s*) is relative to and recessive to wild-type (HD, *s*_*H*_ = 1; SR, *s*_*C*_ =0.05; RD/IRD, *s*_*C*_ =0.2.). We use an initial release of 0.1% HD, and assume recessive lethality of the HD allele and perfect homing (*e*_*HD*_ = *e*_*C*_ = 1). Dashed vertical lines indicate the time of countermeasure release. Large releases (1:1 ratio or countermeasure to pre-countermeasure-release population) are shown in the left column, and small releases (1:10 ratio) are shown in the right column. The split axes with gray bars indicate a change in time scale. **a:** Absent countermeasures, the HD quickly approaches fixation (i.e., would cause population extinction). **b,c:** Release of a SR allele allows a brief increase in HD frequency, followed by a decrease to a low but non-zero equilibrium. **d:** A large RD release yields allelic frequencies after release that are near the stable equilibrium. **e:** A small RD release yields allelic frequencies far from equilibrium, followed by a large transient oscillation, wherein HD frequencies approach zero. **f,g:** Release of IRD results in elimination of HD and wild-type alleles, regardless of release size.

**Figure 2:**
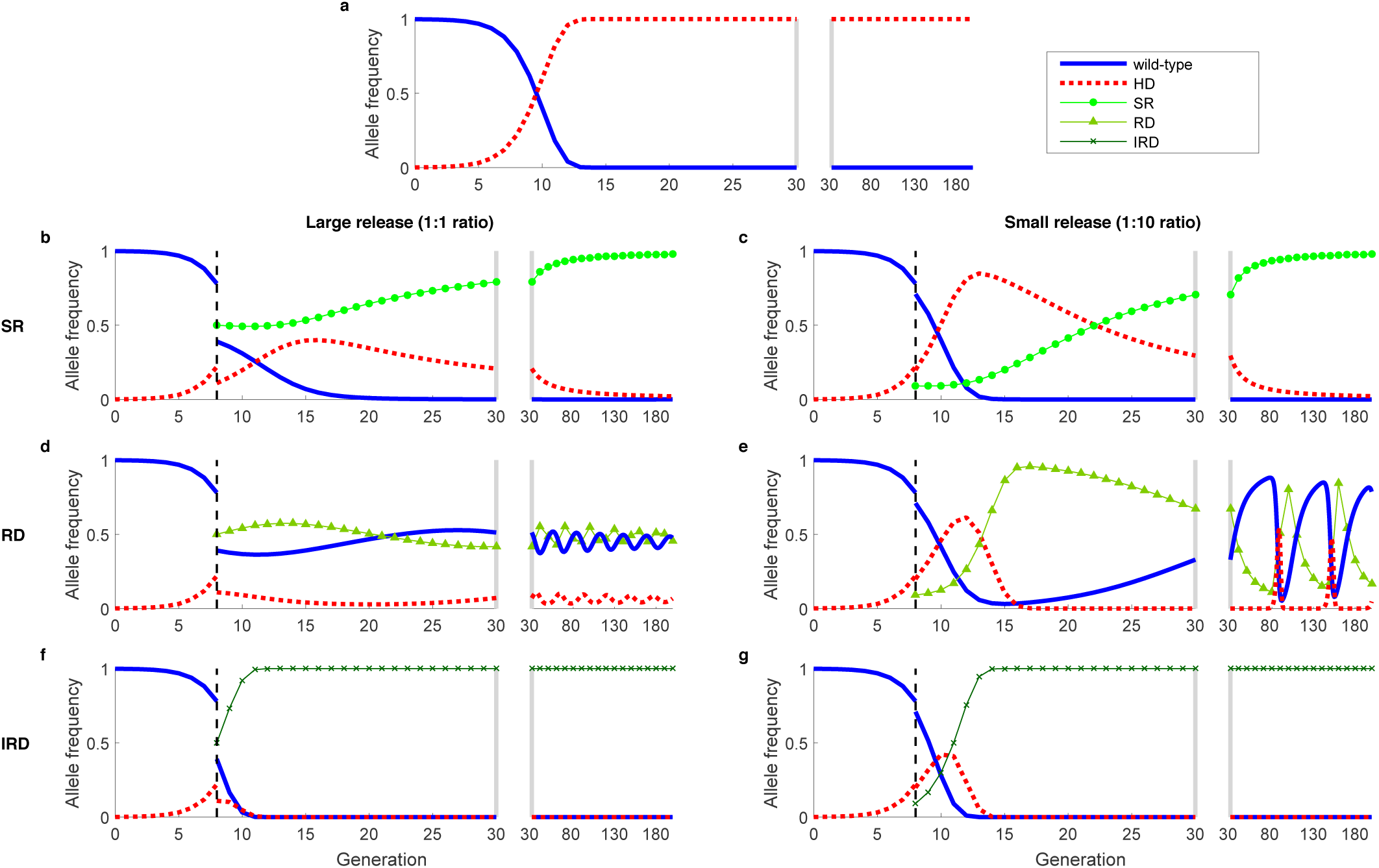
**Dynamics of a replacement HD, alone (a) and with countermeasures (b-g)**, which include a synthetic resistant allele (b,c), reversal drive (d,e), and immunizing reversal drive (f,g). The fitness cost of the HD is *s_H_*=0.3. See Figure 1 for other details. The behavior is qualitatively similar to Figure 1, but the oscillations of the SR and RD are less damped (b-e).

The polymorphic equilibrium and oscillatory dynamics exhibited by the SR and RD systems are due to each alleles frequency-dependent disadvantages relative to other alleles in a rock-paper-scissors type fashion. In this case, the disadvantages result from relative fitness costs and the effects of drive (for the RD), but similar dynamics have been recognized in many unrelated systems[3, 9, 15, 20]. Damped oscillations about a polymorphic equilibrium mean that initial conditions far from the equilibrium result in large fluctuations, temporarily bringing HD frequencies near to zero. Initial conditions close to the equilibrium, on the other hand, do not result in large fluctuations in allelic frequencies, likely allowing the HD to persist (as visualized in a phase plot in Fig. S1a). Initial conditions are not important for determining the fate of the IRD, however, as it does not have frequency-dependent disadvantages to the other alleles.

Relaxing assumptions about fitness but keeping the assumptions of perfect homing and recessive fitness costs in wild-type heterozygotes, we find that a variety of possible stable equilibria may exist for the systems beyond those shown in Figures 1 and 2 (Supplemental Note 1 & Figs. S1-S3). However, given no fitness cost for heterozygotes containing wild-type alleles, a stable, polymorphic equilibrium exists for the SR and RD countermeasures for most plausible combinations of HD, C, and HD/C fitness costs (e.g., when the HD/C heterozygote fitness cost is between the HD and C homozygote fitness costs). Numerically, we find complex eigenvalues of the Jacobian evaluated at the polymorphic equilibrium, which indicate oscillatory dynamics (see Supplemental Note 1). Additionally, assuming additive rather than recessive fitness costs in wild-type heterozygotes changes the regions of parameter space that result in each equilibrium for the SR and RD countermeasures but still results in uncertain removal of the HD for SR and RDs, and likely removal of the HD for IRDs (Figs. S4-S5).

With a deterministic model, likelihood of stochastic extinction during transient oscillations cannot be measured directly, but the likelihood increases as the minimum frequency decreases. Figures 3 and 4 show the minimum HD frequency achieved within the first 100 generations after countermeasure release for varying fitness costs and initial conditions, returning to the assumption of recessive fitness costs and that the cost to the HD/C heterozygote is the minimum of the HD and C fitness costs. In general, low countermeasure fitness costs yield the greatest reductions in HD frequencies, both for RD (Fig. 3) and for SR (Fig. 4) countermeasures, by lowering the HD frequency at the polymorphic equilibrium. When HD frequencies are low, an RD released in numbers close to that of the current population (1:1 ratio) causes the system to quickly approach the polymorphic equilibrium instead of exhibiting large transient oscillations that bring the HD frequency near to 0. A very large RD release that immediately limits HD/wild-type mating would likely cause stochastic HD elimination (bottom row) in a randomly mating population, but would require additional time and resources necessary to rear and release a sufficient number of RD individuals. Unlike the RD, SR cannot be effective when its fitness cost exceeds that of the HD (Fig. 4, top-left corner of each panel). As with the RD, SR releases in size equal to the pre-release population bring the system near to its polymorphic equilibrium, which would prevent the HD frequency from transiently reaching very small frequencies. However, because oscillations occur on a slower time-scale than with the RD (see Fig. 1b/e), the minimum is not always reached within 100 generations. For an IRD, all panels reach minimum frequencies by generation 100.

**Figure 3:**
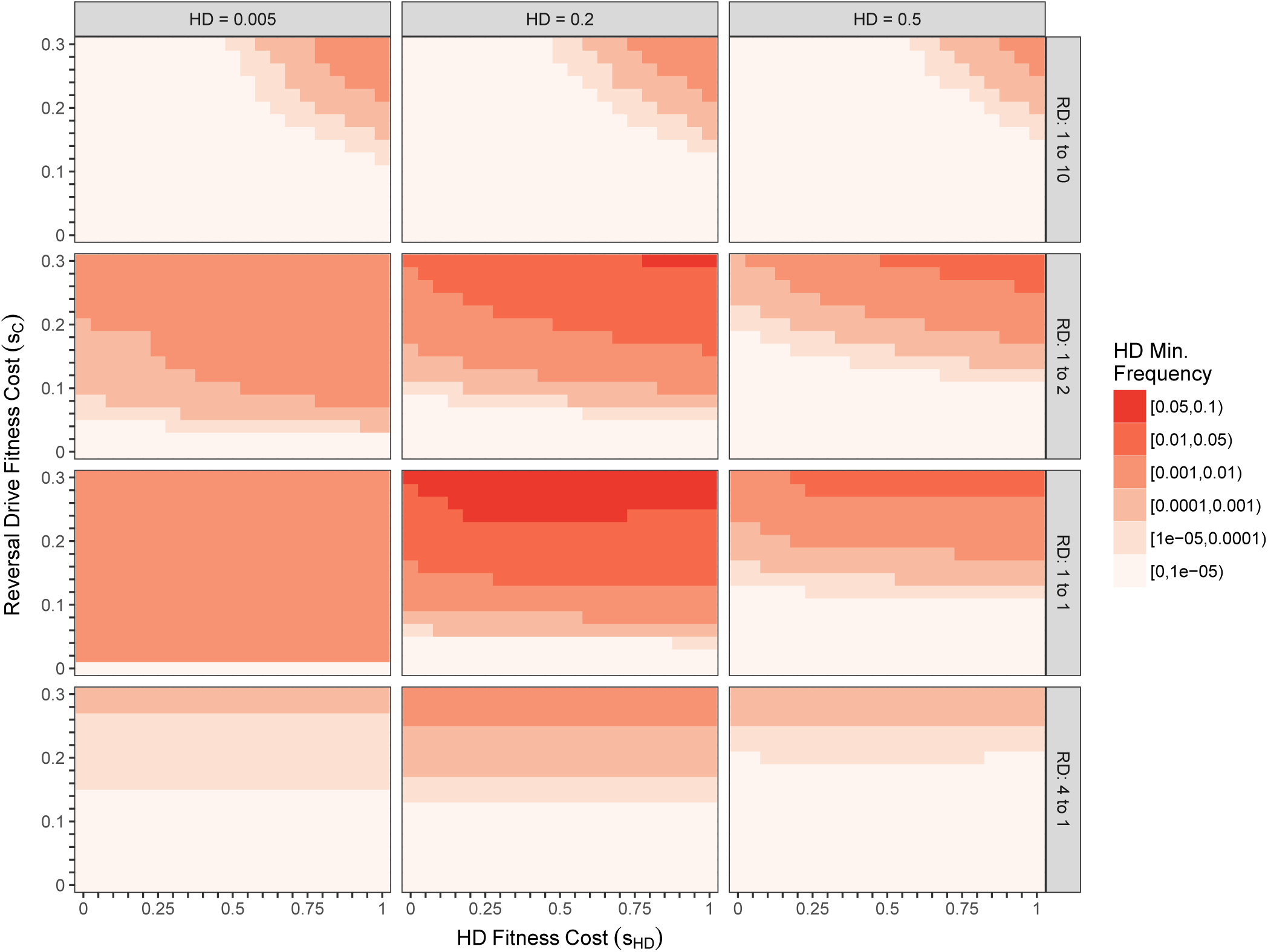
Minimum HD allele frequency in the first 100 generations after RD release for various fitness costs, initial conditions, and release ratios. Light shades indicate higher likelihood of stochastic loss of HD, while dark shades highlight instances where removal of the HD is less likely. Axes show fitness costs of the HD (x-axis) and RD (y-axis). Initial conditions vary between panels: columns vary the HD pre-release frequency, and rows vary the RD release size, which is shown as a release ratio (e.g., 4 to 1 releases 4 RD alleles for every pre-release allele). We assume recessive fitness costs and perfect homing. The largest HD fitness cost (*s*_*HD*_ = 1) corresponds to a suppression HD, whereas small HD fitness costs correspond to a replacement HD. Note that maximum HD frequency varies independently from minimum HD frequency; in small RD releases (top row of panels), the HD frequency can experience large increases before dropping to the low minimum levels show here. Overall, a RD release appears least likely to eliminate a target HD when RD fitness costs are large, and when the RD release yields post-release frequencies near the equilibrium. The higher minimum frequency for larger HD fitness costs in many panels is due to the smaller amplitudes of oscillations compared to systems with lower HD fitness costs, as seen Figures 1 and 2. Smaller oscillations result in the system tending directly toward the equilibrium.

**Figure 4:**
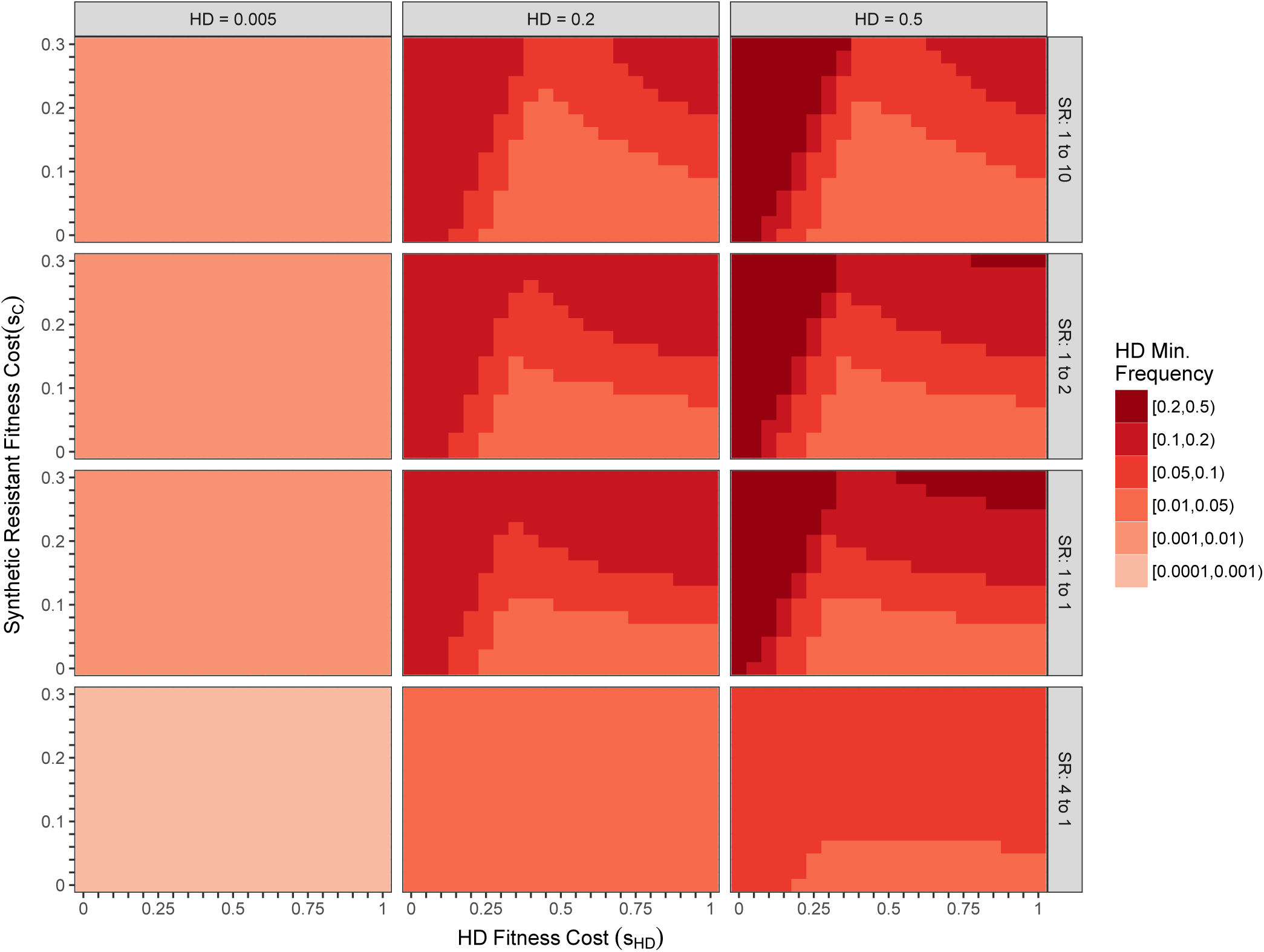
Minimum HD allele frequency in the first 100 generations after SR release for various fitness costs, initial conditions, and release ratios. See Figure 3 for details, noting that the legend colors refer to different minimum frequencies. Similarly to the RD, the SR is least likely to eliminate a target HD when its fitness costs are large, and when the release yields post-release frequencies near the equilibrium, though equilibrium frequencies are not identical to RDs. In some of the simulations, the system is not yet at equilibrium, and the HD is still decreasing in frequency at 100 generations.

Finally, we further relax our assumptions to account for less than perfect homing with the creation of naturally resistant alleles (Fig. 5). The qualitative behavior found in the case of perfect homing remains, except that the IRD eventually falls out of the population since it has lower fitness than naturally resistant alleles. Given imperfect homing, HD frequencies would fall even in the absence of countermeasures. As with SR, though, the HD is sustained long-term due to a stable, polymorphic equilibrium (Fig. 5a).

**Figure 5:**
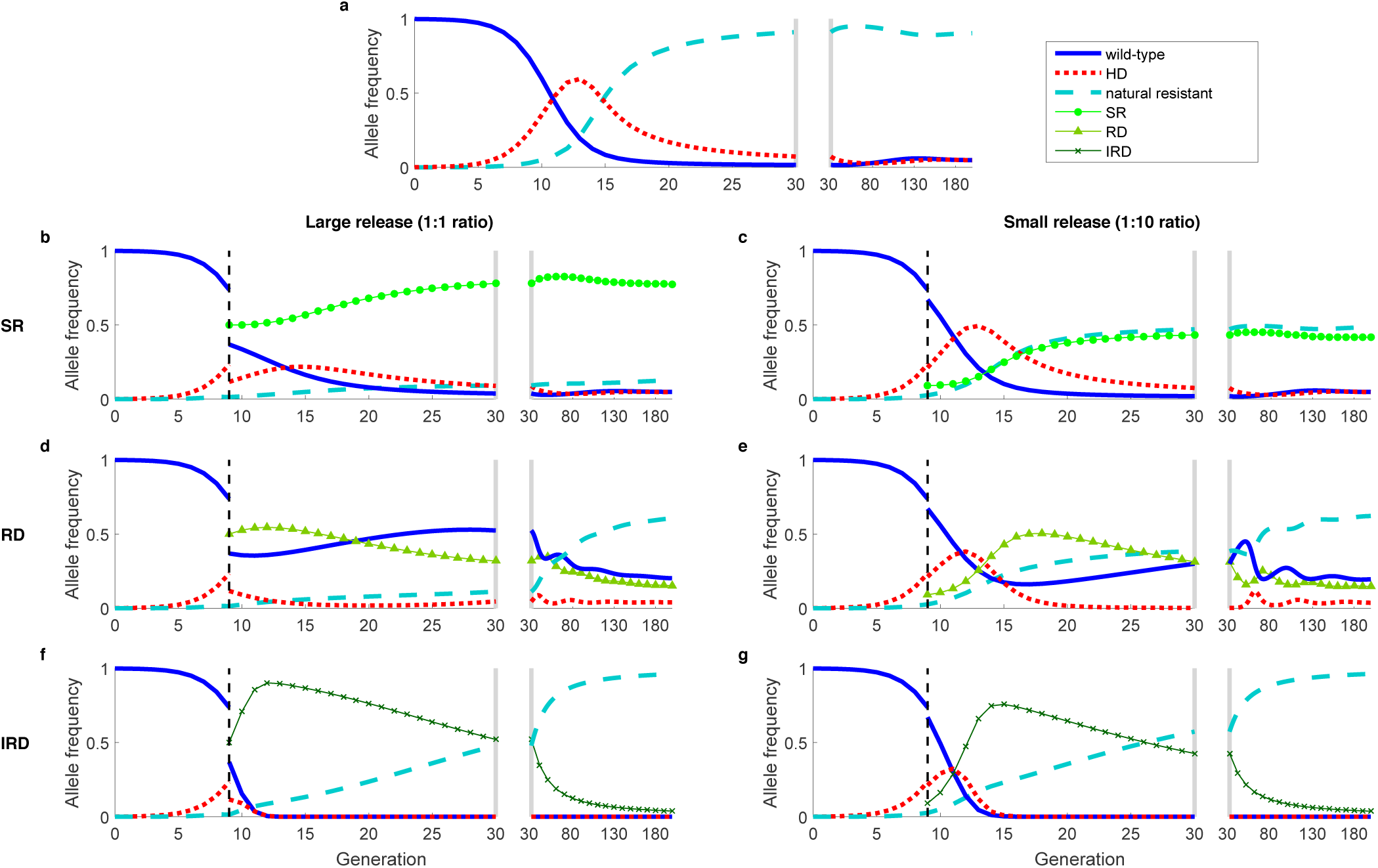
**Dynamics of an imperfect suppression HD, where the drive fails and produces naturally resistant alleles, alone (a) and with countermeasures (b-g)**, which include a synthetic resistant allele (b,c), reversal drive (d,e), and immunizing reversal drive (f,g). Homing is imperfect (*e*_*H*_ = *e*_*C*_ = 0.9), and unsuccessful homing results in natural resistance via NHEJ with fitness cost *s*_*R*_=0.05. See Figure 1 for other details. The biggest change from accounting for imperfect homing is that the IRD falls out of the population in the long-term (f,g) because of low-fitness cost alleles resistant to cutting.

## Discussion

A variety of genetic approaches have been proposed to counter unintended effects of an HD, but there has been limited theoretical evaluation of these approaches. Here we compare the dynamics of SR, RD, and IRD countermeasures upon release into a population prior to HD fixation and find that the long-term behavior of the system differs greatly between countermeasures. In particular, SR and RD countermeasures are not guaranteed to eliminate an HD from a population because these systems often exhibit a stable polymorphic equilibrium. Elimination of the HD via SR or RD becomes less likely with higher countermeasure fitness costs, as the equilibrium HD frequency is further from zero, and oscillations around the polymorphic equilibrium are less likely to cause stochastic loss of the HD. Due to the small magnitude of oscillations with release conditions close to equilibrium, the frequencies of the HD prior to release and the relative size of the countermeasure release are important factors in determining the likelihood of HD elimination. If either of these countermeasures were to fully eliminate the HD, the wild-type allele would ultimately recover to fixation as long its fitness is higher than the countermeasure.

An IRD that targets both HD and wild-type alleles, on the other hand, would theoretically ensure the rapid removal of the HD from the population, but would also result in the Cas9 gene and guide RNAs remaining in the population. Implications of leaving Cas9 in the population are unclear, such as the likelihood of off-target effects, and future research should seek to evaluate such effects. If any naturally resistant alleles develop, or with the release of an SR allele, the IRD would eventually fall out of the population, provided that the cost of the IRD is greater than the resistant allele. These qualitative differences between countermeasures must be considered when deciding whether they are suitable tools for mitigating adverse effects of an HD.

The model and subsequent analysis presented here yields critical insights into the qualitative behavior of, and differences between, genetic countermeasures. Nonetheless, future work could explore several additional aspects of HD-based countermeasures and provide quantitative risks associated with them. Models that track population size as well as allele frequency, and that incorporate demographic stochasticity, could be used to better assess options for eliminating suppression HDs. For suppression HDs, population size could drastically decrease, and the effects of genetic drift could predominate[19]. Also deserving of increased attention are the effects of spatial heterogeneity. In particular, spatial isolation of small populations could limit an alleles spread, potentially impacting countermeasure success. Incorporating spatial heterogeneity could also be useful in assessing the impact of movement between the target population and nearby populations on the long-term fates of the relevant constructs. Important consequences of movement include the likelihood of HD spillover to nearby populations, and whether immigration of wild-type organisms could sustain a HD in a system where stochastic elimination is otherwise likely. Effects of spatial heterogeneity may be different for RDs and IRDs, so follow-up modeling studies will be needed. Finally, effects of assumptions about natural resistance to homing drives should be explored further. While some work has explored the development of natural resistance to HDs[1, 2, 5, 16, 18, 21], these findings should be updated as HD limitations are understood.

Many have proposed countermeasures as emergency tools to mitigate unintended negative effects that might arise after release of an HD. However, to date only limited theoretical analysis has addressed countermeasures’ abilities to reverse HDs. Additionally, discussion about countermeasures has often been ambiguous regarding differences between types of countermeasures and expectations of countermeasure outcomes. Depending on the severity of unintended effects, countermeasures may have the goal of simply halting the spread of an HD, or possibly removing an HD from the population and returning the population to its original state. This work is motivated by a desire to more clearly specify differences between various countermeasure strategies, as well as to critically assess potential outcomes. Here we show that the RD does not eliminate the HD for certain release conditions and fitness parameters. The existence of a polymorphic equilibrium with oscillatory dynamics allows for the HD allele frequency to initially increase, to remain constant, or to decrease, depending on the reversal release size. In such cases, larger countermeasure fitness costs decrease the likelihood of long-term eradication of the HD allele. IRDs are expected to effectively eliminate the HD in a timely manner but leave Cas9 present in the population, though any resistant alleles would cause the IRD to eventually fall out of the population. RDs leave only guide RNAs if they successfully eliminate the HD, but given any fitness cost to the RD, the wild-type would be expected to return. Overall, these results show that no single countermeasure, as currently proposed, should be considered a silver bullet for mitigating unintended effects of HDs. As such, we recommend careful examination of risks associated with each of the countermeasures limitations prior to release.

## Acknowledgements

This research was funded by the National Institutes of Health (NIH) grant R01-AI091980, the W. M. Keck Foundation, and the National Science Foundation (RTG/DMS - 1246991 and NSF-IGERT - 1068676). We thank Brandon Hollingsworth, Jaye Sudweeks, Sumit Dhole, Jennifer Baltzegar, Kevin Esvelt, Ethan Bier, Anthony James, and Valentino Gantz for helpful discussion.

## Author contributions statement

All authors conceptualized the research, MRV developed the mathematical model and wrote the initial draft, and all authors edited, revised, and reviewed the manuscript.

## Additional information

### Data availability

Model code can be requested from authors.

## References

[1] Bull, J. J. (2016). Oup: lethal gene drive selects inbreeding. Evolution, Medicine, and Public Health, 2017(1):1–16.

[2] Burt, A. (2003). Site-specific selfish genes as tools for the control and genetic engineering of natural populations. Proceedings of the Royal Society of London B: Biological Sciences, 270(1518):921–928.

[3] Cameron, D. D., White, A., and Antonovics, J. (2009). Parasite–grass–forb interactions and rock–paper–scissor dynamics: predicting the effects of the parasitic plant rhinanthus minor on host plant communities. Journal of Ecology, 97(6):1311–1319.

[4] Champer, J., Buchman, A., and Akbari, O. S. (2016). Cheating evolution: engineering gene drives to manipulate the fate of wild populations. Nature Reviews Genetics, 17(3):146–159.

[5] Champer, J., Reeves, R., Oh, S. Y., Liu, C., Liu, J., Clark, A. G., and Messer, P. W. (2017). Novel crispr/cas9 gene drive constructs in drosophila reveal insights into mechanisms of resistance allele formation and drive efficiency in genetically diverse populations. bioRxiv, page 112011.

[6] Deredec, A., Burt, A., and Godfray, H. C. J. (2008). The population genetics of using homing endonuclease genes in vector and pest management. Genetics, 179(4):2013–2026.

[7] DiCarlo, J. E., Chavez, A., Dietz, S. L., Esvelt, K. M., and Church, G. M. (2015). Safeguarding crispr-cas9 gene drives in yeast. Nature biotechnology, 33(12):1250–1255.

[8] Doudna, J. A. and Charpentier, E. (2014). The new frontier of genome engineering with crispr-cas9. Science, 346(6213):1258096.

[9] Durrett, R. and Levin, S. (1997). Allelopathy in spatially distributed populations. Journal of theoretical biology, 185(2):165–171.

[10] Esvelt, K. M., Smidler, A. L., Catteruccia, F., and Church, G. M. (2014). Concerning rna-guided gene drives for the alteration of wild populations. Elife, 3:e03401.

[11] Gantz, V. M. and Bier, E. (2015). The mutagenic chain reaction: A method for converting heterozygous to homozygous mutations. Science, 348(6233):442–444.

[12] Gantz, V. M. and Bier, E. (2016). The dawn of active genetics. BioEssays, 38(1):50–63.

[13] Hartl, D. and Clark, A. (2007). Principles of population genetics. sunderland (ma).

[14] Hsu, P. D., Lander, E. S., and Zhang, F. (2014). Development and applications of crispr-cas9 for genome engineering. Cell, 157(6):1262–1278.

[15] Kerr, B., Riley, M. A., Feldman, M. W., and Bohannan, B. J. (2002). Local dispersal promotes biodiversity in a real-life game of rock–paper–scissors. Nature, 418(6894):171–174.

[16] Marshall, J. M., Buchman, A., et al. (2017). Overcoming evolved resistance to population-suppressing homing-based gene drives. Scientific Reports, 7.

[17] National Academies of Sciences, Engineering, and Medicine (2016). Gene Drives on the Horizon: Advancing Science, Navigating Uncertainty, and Aligning Research with Public Values. National Academies Press.

[18] Noble, C., Olejarz, J., Esvelt, K. M., Church, G. M., and Nowak, M. A. (2017). Evolutionary dynamics of crispr gene drives. Science Advances, 3(4):e1601964.

[19] Okamoto, K. W., Robert, M. A., Gould, F., and Lloyd, A. L. (2014). Feasible introgression of an anti-pathogen transgene into an urban mosquito population without using gene-drive. PLoS Negl Trop Dis, 8(7):e2827.

[20] Sinervo, B. and Lively, C. M. (1996). The rock-paper-scissors game and the evolution of alternative male strategies. Nature, 380(6571):240.

[21] Unckless, R. L., Clark, A. G., and Messer, P. W. (2017). Evolution of resistance against crispr/cas9 gene drive. Genetics, 205(2):827–841.

[22] Unckless, R. L., Messer, P. W., Connallon, T., and Clark, A. G. (2015). Modeling the manipulation of natural populations by the mutagenic chain reaction. Genetics, pages genetics-115.

[23] Wu, B., Luo, L., and Gao, X. J. (2016). Cas9-triggered chain ablation of cas9 as a gene drive brake. Nature biotechnology, 34(2):137–138.

[24] Zetsche, B., Gootenberg, J. S., Abudayyeh, O. O., Slaymaker, I. M., Makarova, K. S., Essletzbichler, P., Volz, S. E., Joung, J., van der Oost, J., Regev, A., et al. (2015). Cpf1 is a single rna-guided endonuclease of a class 2 crispr-cas system. Cell, 163(3):759–771.

